# Proteome-by-phenome Mendelian Randomisation detects 38 proteins with causal roles in human diseases and traits

**DOI:** 10.1101/631747

**Authors:** Andrew D. Bretherick, Oriol Canela-Xandri, Peter K. Joshi, David W. Clark, Konrad Rawlik, Thibaud S. Boutin, Yanni Zeng, Carmen Amador, Pau Navarro, Igor Rudan, Alan F. Wright, Harry Campbell, Veronique Vitart, Caroline Hayward, James F. Wilson, Albert Tenesa, Chris P. Ponting, J. Kenneth Baillie, Chris Haley

## Abstract

Target identification remains a crucial challenge in drug development. To enable unbiased detection of proteins and pathways that have a causal role in disease pathogenesis or progression, we propose proteome-by-phenome Mendelian Randomisation (P^2^MR). We first detected genetic variants associated with plasma concentration of 249 proteins. We then used 64 replicated variants in two-sample Mendelian Randomisation to quantify evidence of a causal role for each protein across 846 phenotypes: this yielded 509 robust protein-outcome links. P^2^MR provides substantial promise for drug target prioritisation. We provide confirmatory evidence for a causal role for the proteins encoded at multiple cardiovascular disease risk loci (*FGF5, IL6R, LPL, LTA*), and discovered that intestinal fatty acid binding protein (*FABP2*) contributes to disease pathogenesis. Additionally, we find and replicate evidence for a causal role of tyrosine-protein phosphatase non-receptor type substrate 1 (SHPS1; *SIRPA*) in schizophrenia. Our results provide specific prediction of the effects of changes of plasma protein concentration on complex phenotypes in humans.

---

An initial goal of drug development is the identification of targets – in most cases, proteins – whose interaction with a drug ameliorates the development, progression, or symptoms of disease. After some success, the rate of discovery of new targets has not accelerated despite substantially increased investment (Munos, 2009). A large proportion of drugs fail at the last stages of development – clinical trials – because their targets do not alter whole-organism phenotypes as expected from pre-clinical research (Arrowsmith, 2011).

Preclinical science is engaging with increasingly complex systems in which prediction of the effects of an intervention is ever more difficult (Civelek & Lusis, 2014). The ability to cut through complexity to distinguish factors that modulate whole-organism phenotypes is a major advantage of genetic (Baillie, 2014) and functional genomic (Baillie et al., 2018) approaches to drug development. Nevertheless, genetic associations with disease are not immediately interpretable (MacArthur et al., 2017): most disease-associated variants fail to alter protein-coding sequence, but instead alter protein levels via often poorly understood molecular mechanisms.

A subset of disease states have been studied with adequately-powered genome-wide association (GWA) studies (Finan et al., 2017). From these, persuasive evidence already exists for the utility of using genetic and genomic techniques to inform drug development: the presence of genetic evidence in support of a protein could double the probability of success in clinical trials for drugs targeting that protein (M. R. Nelson et al., 2015). In a recent study, 12% of all targets for licenced drugs could be rediscovered using GWA studies (Finan et al., 2017). However, these GWA study approaches generally rely on measures of proximity of a disease-associated genetic variant to a protein-coding gene, and proximity alone does not imply causality.

Mendelian Randomisation (MR) uses genetic variants to provide an estimate of the effect of an exposure on an outcome, using the randomness of assignment of genotype to remove the effects of unmeasured confounding (Smith & Ebrahim, 2003). The approach is analogous to a naturally-occurring randomised control trial. When a genetic variant predicts the abundance of a mediator, MR tests the hypothesis that this mediator plays a causal role in disease risk. This is possible because the patient or participant was effectively randomised at conception to a genetically-determined level of that mediator. Under this model, it is possible to use population level genetic information to draw causal inference from observational data. However, there are, as with any study, unverifiable assumptions with this study design: a major concern is that alternative causal pathways may link the instrumental variable (here, the DNA variant) to the phenotype (the disease outcome). In a clinical trial this would be analogous to a drug influencing a disease through a different pathway than via its reported target. In MR, addressing the risk of alternative causal pathways is of great practical importance in order to avoid pursuing drugs that target an irrelevant molecular entity, and hence that have no beneficial effect. In order to address this, we limited ourselves to using locally-acting pQTLs as instrumental variables. We believe this approach provides stronger supporting evidence for causation than relying on proximity of a disease-associated genetic variant to a gene, or using mRNA abundance as a proxy for protein abundance (Mirauta et al., 2018).

Due to recent advances in proteomic technologies, the availability of pQTL data has increased dramatically in recent years (Folkersen et al., 2017; Suhre et al., 2017; Sun et al., 2018; Yao et al., 2018). A number of these studies attempt to infer causality using MR and similar techniques. In our approach, we applied pQTL based MR in a data-driven manner across the full range of phenotypes available in GeneAtlas (UK Biobank (Canela-Xandri, Rawlik, & Tenesa, 2017)), as well as supplementing this with additional studies identified through Phenoscanner (Staley et al., 2016). We performed GWA for 249 proteins in two European cohorts, and then adopted a proteome-by-phenome Mendelian randomisation (P^2^MR) approach to assess the potential causal role of 64 proteins in 846 outcomes (e.g. diseases, anthropomorphic measures, etc.). GeneAtlas results were further stratified according to their consistency with a single underlying causal variant (affecting both variation in protein concentration and outcome phenotype) or otherwise. Ultimately, of the 249 proteins, 38 were identified as causally contributing to human disease or other quantitative trait.

## Results

The abundance of an individual protein may be associated with DNA variants both local and distant to its gene (termed local- and distal-pQTLs, respectively). We assayed the plasma levels of 249 proteins using high-throughput, multiplex immunoassays and then performed genome-wide association of these levels in two independent cohorts (discovery and replication) of 909 and 998 European individuals. P^2^MR was applied to 54,144 exposure-outcome pairs obtained from 64 significantly (p-value <5×10^−8^) associated, replicated (Bonferroni correction for multiple testing), local-pQTLs, and 778 phenotypes obtained from GeneAtlas (UK Biobank (Canela-Xandri et al., 2017)) and 68 phenotypes from 20 additional genome-wide association (meta-analysis) studies (The CARDIoGRAMplusC4D Consortium et al., 2015; R. A. Scott et al., 2017; C. P. Nelson et al., 2017; Liu et al., 2015; Schizophrenia Working Group of the Psychiatric Genomics Consortium et al., 2014; Bronson et al., 2016; Okada et al., 2014; van Rheenen et al., 2016; Hammerschlag et al., 2017; Sniekers et al., 2017; Okbay et al., 2016; Hou et al., 2016; Beaumont et al., 2018; Phelan et al., 2017; van der Harst & Verweij, 2018; Berg et al., 2016; de Moor et al., 2015; The EArly Genetics and Lifecourse Epidemiology (EAGLE) Eczema Consortium et al., 2015; M. A. Ferreira et al., 2017; Astle et al., 2016) identified through Phenoscanner (Staley et al., 2016) (Figure 1; Supplementary Table S1; Methods).

**Figure 1.**
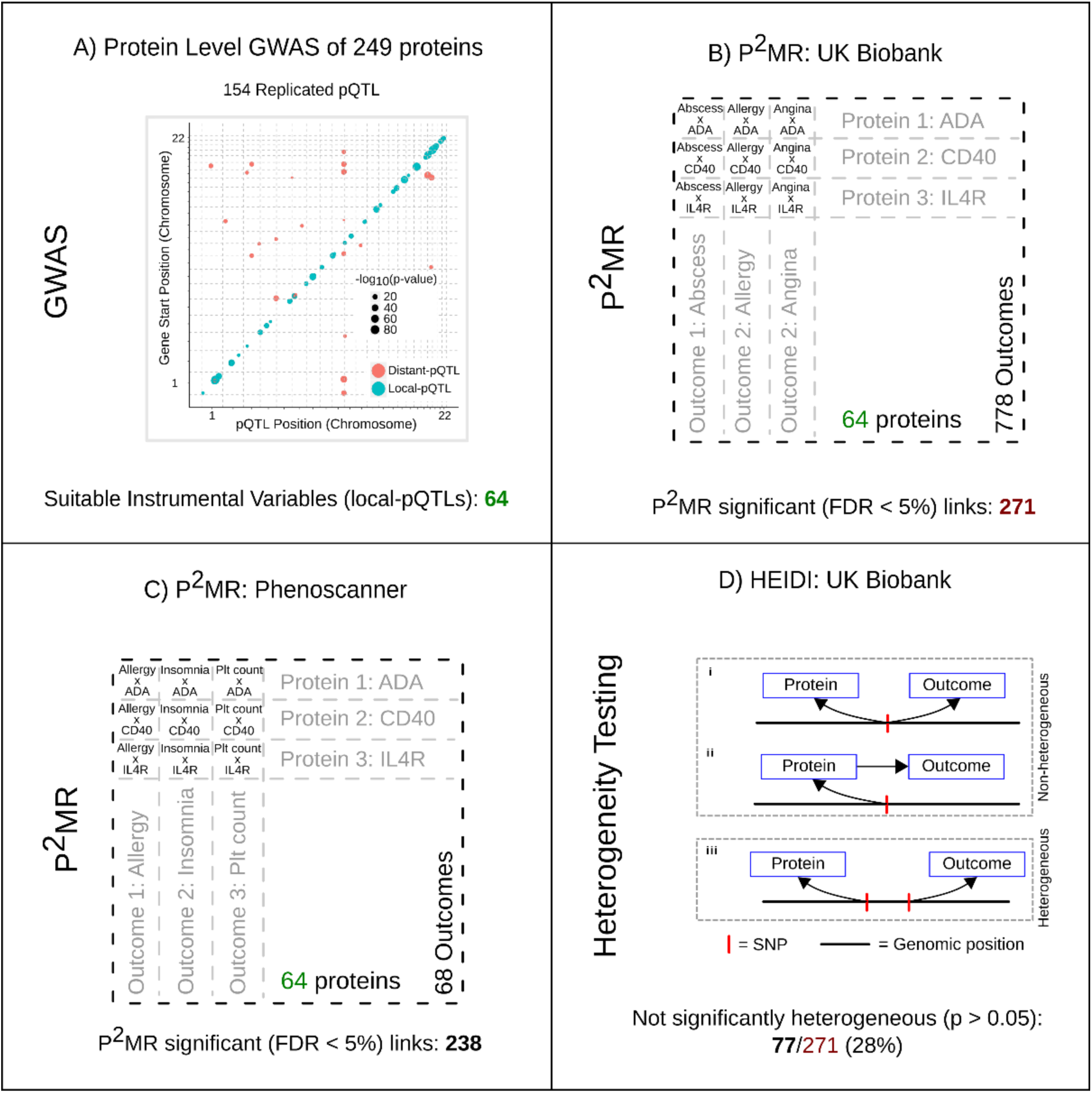
Proteome-by-phenome Mendelian Randomisation (P^2^MR). A) Genome-wide associations of the plasma concentrations of 249 proteins from two independent European cohorts (discovery and replication) were calculated. The plot shows pQTL position against chromosomal location of the gene that encodes the protein under study for all replicated pQTLs. The area of a filled circle is proportional to its - log10(p-value) in the replication cohort. Blue circles indicate pQTLs ±150kb of the gene (‘local-pQTLs’); red circles indicate pQTLs more than 150kb from the gene. B, C) Local-pQTLs of 64 proteins were taken forward for P^2^MR analysis. These were assessed against 778 outcome phenotypes from GeneAtlas (Canela-Xandri et al., 2017) (panel B; UK Biobank) and 68 phenotypes identified using Phenoscanner (Staley et al., 2016) (panel C). In each set of results an FDR of <0.05 was considered significant. D) Heterogeneity in dependent instruments (HEIDI (Zhu et al., 2016)) testing was undertaken for MR significant results from UK Biobank (n = 271). This test seeks to distinguish a single causal variant at a locus effecting both exposure and outcome directly (as in i) or in a causal chain (as in ii), from two causal variants in linkage disequilibrium (as in iii), one effecting the exposure and the other effecting the outcome.

In total, we identified 509 protein-outcome links for which there is evidence of a causal role of the exposure (protein) on the outcome phenotype (trait).

### pQTLs

pQTLs were highly concordant between the two cohorts (Supplementary Table S2). Of the 209 independent pQTLs identified in the discovery cohort (p-value <5×10^−8^), 154 were successfully replicated (Bonferroni correction for multiple testing; consistent direction of effect). These represented pQTLs for 82 proteins, all but two encoded by autosomal genes. Lead variants (smallest p-value within the locus; Methods) were identified at each locus, the majority (64/80; 80%) of these proteins had the lead variant of one or more pQTLs located close to the gene encoding the protein (±150kb; Figure 1) and hence were used as instrumental variables suitable for MR. In many respects, locally-acting pQTLs are ideal instrumental variables: they have large effect sizes, have highly plausible biological relationships with protein level, and provide quantitative information about (often) directly druggable protein targets. This is in contrast to distal pQTLs: the pathway through which they exert their effects is generally unknown, with no *a priori* expectation of a direct effect on a single target gene.

### Outcome GWA Studies

Results linking the genetic variants and outcome traits and diseases were obtained from secondary cohorts. UK Biobank has captured a wealth of information on a large – approximately 500,000 individuals – population cohort that includes anthropometry, haematological traits, and disease outcomes. Genome-wide association of 778 phenotypes from UK Biobank has been performed and published as GeneAtlas (Canela-Xandri et al., 2017). Although the cohort is large, for many diseases the number of UK Biobank individuals affected is small, resulting in low statistical power. Consequently, we augmented these results with additional studies identified using Phenoscanner (Staley et al., 2016) (Methods).

### Mendelian Randomisation

MR depends upon an assumption that the DNA variant used as an instrumental variable is robustly associated with the exposure. In our case we ensured this by using stringent discovery and replication criteria for instrument selection. By limiting ourselves to using locally-acting pQTLs as instruments, we sought to leverage *a priori* biological knowledge regarding cellular protein production to substantially increase confidence in the existence of a direct path from DNA variant to protein, and from protein to outcome. P^2^MR yielded 271 protein-outcome pairs that were significant (false discovery rate (FDR) <0.05) in UK Biobank, and 238 significant (FDR <0.05) pairs using data from Phenoscanner. Thirty-two of the 64 proteins were causally implicated for one or more outcomes in UK Biobank, and 36 of 64 in the outcomes identified through Phenoscanner studies. The outcomes from GeneAtlas and Phenoscanner are not mutually exclusive, and some of the studies included from Phenoscanner included data from UK Biobank, however, overall, 38 of the 64 proteins (60%) were implicated in at least one outcome (Supplementary Tables S3 and S4).

Proteins were implicated in diseases ranging from schizophrenia to cardiovascular disease (Figure 2, Supplementary Tables S3 and S4). We applied a method, HEIDI (Zhu et al., 2016), which explicitly accounts for the linkage disequilibrium (LD) structure of the locus to assess the heterogeneity of MR effect estimates between the lead variant (the primary instrument) and those of linked variants. HEIDI tests the hypothesis that the observed MR results are caused by two distinct causal variants. Of the UK Biobank causal inferences, 77 survived the HEIDI heterogeneity test (p-value >0.05). Therefore, these 77 proteins have (a) high-quality evidence of association to a DNA variant which provides congruent predictions for both plasma protein levels and disease risk / outcome phenotype, and (b) because of the physical proximity to the SNP to the coding-sequence of the gene for the protein, and non-significant HEIDI result, a low risk of pleiotropy (Supplementary Tables S3). These pairs thus provide the most robust evidence that the level of the protein directly alters disease risk / outcome phenotype.

**Figure 2.**
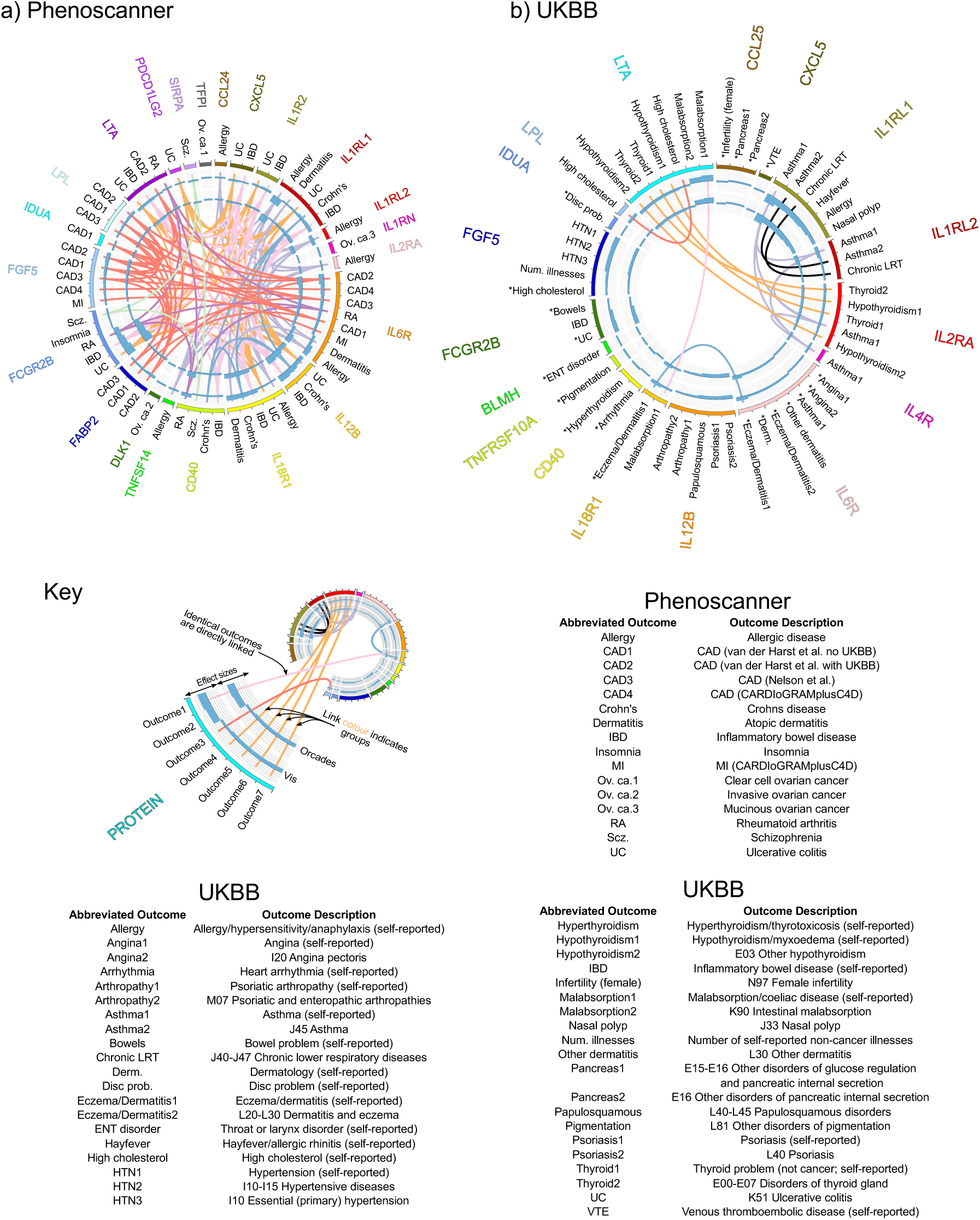
Significant (FDR <0.05) P^2^MR protein-outcome causal inferences: disease subset. a) Phenoscanner (Staley et al., 2016): P^2^MR significant protein-disease outcome causal inferences for 20 Phenoscanner studies. b) GeneAtlas (Canela-Xandri et al., 2017): MR significant protein-disease outcome causal inferences for UK Biobank data. Asterisks indicate P^2^MR estimates that are not significantly heterogeneous (HEIDI, Main Text (Zhu et al., 2016)). Graphical key: Reading from the outside in: protein (exposure; HGNC symbol); disease outcome; key colour; bar chart of the signed (beta/standard error)^2^ value of the MR estimate (using pQTL data from the discovery cohort; Methods); and bar chart of the signed (beta/standard error)^2^ value of the MR estimate (using pQTL data from the replication cohort; Methods). Central chords join identical outcomes. Identically coloured chords indicate similar outcome groups, e.g. thyroid disease.

**Figure 3:**
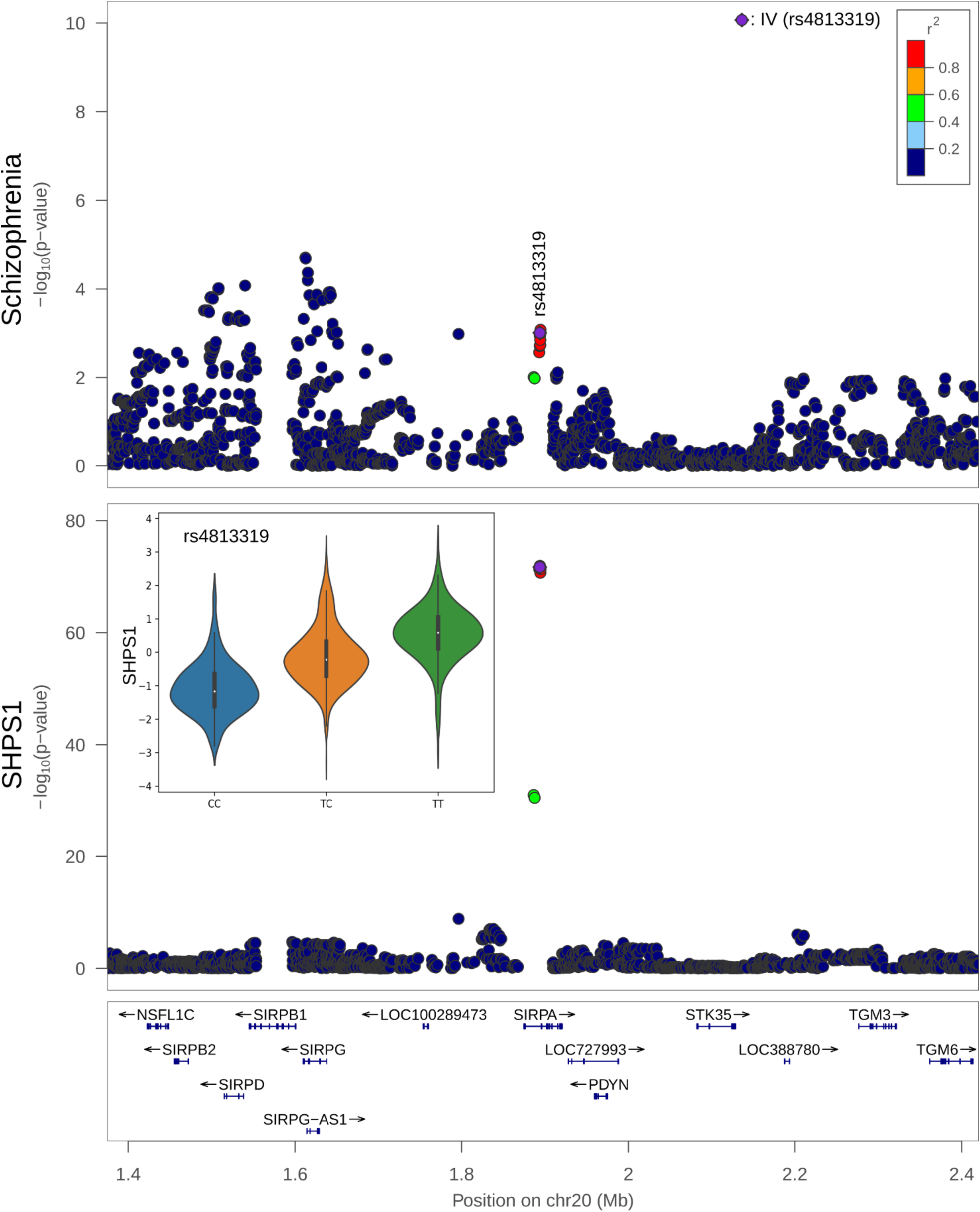
Co-localisation of SHPS1 (encoded by *SHPS1*: synonym *SIRPA*) and schizophrenia DNA associations. Upper panel, locuszoom (Pruim et al., 2010) of the region surrounding *SHPS1* and the associations with schizophrenia (Schizophrenia Working Group of the Psychiatric Genomics Consortium et al., 2014); lower panel, associations with SHPS1. Lower panel inset, the relative concentration of SHPS1 across the 3 genotypes of rs4813319 – the DNA variant used as the instrumental variable (IV) in the MR analysis: CC, CT, and TT.

However, all 509 causal inferences (271 from GeneAtlas (Canela-Xandri et al., 2017) and 238 from studies identified through Phenoscanner (Staley et al., 2016); Figures 2, S1, S2, S3, S4, and Tables S3 and S4), even those consistent with heterogeneity, remain potential high quality drug targets. This is because the HEIDI heterogeneity test (Figure 1) is susceptible to type I errors in this context, as it does not account for multiple causal variants in a locus. In addition, we apply HEIDI in a conservative manner: as a significant HEIDI test implies heterogeneity, we did not apply a multiple testing correction. If a Bonferroni correction (271 tests) were to be applied to the HEIDI p-value, 180 of the protein-outcome pairs are not significantly heterogeneous.

For some of these inferences, genetic evidence of an association between a protein and phenotype has been proposed based on physical proximity of the genes to GWA intervals. For nearly two-thirds (62%; 318/509) however, significant (FDR <0.05) MR association between protein and outcome was not matched by significant (p-value <5×10^−8^) association of the DNA variant to outcome. This suggests that P^2^MR has a greater potential to link protein product and phenotype than naïve genome-wide association.

Our results draw causal inference between protein concentration and disease, for example, IL4R and asthma, IL2RA and thyroid dysfunction, and IL12B and psoriasis (Figure 2). Taking IL6R as an example, we found evidence for a causal association between plasma IL6R abundance and coronary artery disease (CAD), atopy, and rheumatoid arthritis (Figure 2, and Tables S3 and S4). We note that: 1) tocilizumab (an IL6 receptor antagonist) is in clinical use for treating rheumatoid arthritis (L. J. Scott, 2017), 2) there is prior evidence from MR demonstrating elevated levels of soluble IL6R and reduced cardiovascular disease (IL6R Genetics Consortium Emerging Risk Factors Collaboration, 2012; Interleukin-6 Receptor Mendelian Randomisation Analysis (IL6R MR) Consortium et al., 2012), and 3) the evidence of a causal link between IL6R and atopy was not well established previously. Notably however, tocilizumab has been used to treat three atopic dermatitis patients, and all patients experienced >50% improvement in disease (Navarini, French, & Hofbauer, 2011). In addition, Ullah et al. (2014) demonstrated that tocilizumab caused a reduction in Th2/Th17 response and associated airway inflammatory infiltration in a mouse model of experimental allergic asthma.

As further illustration, we take two clinically important phenotypes as case-studies: CAD risk and schizophrenia.

### CAD and FABP2

P^2^MR identified 5 proteins as contributing to CAD pathogenesis: FABP2, FGF5, IL6R, LPL, and LTA. Of these, 4 (FGF5, LPL, IL6R, and LTA) had been implicated previously (Klarin et al., 2017; C. P. Nelson et al., 2017; Ozaki et al., 2002), whereas FABP2 had more limited evidence for its involvement.

FABP2 (intestinal fatty acid-binding protein) is causally linked by P^2^MR to CAD (Figure 2). A FABP2 non-synonymous mutation (Ala54Thr) had been proposed as a risk factor for CAD (Yuan, Yu, & Zeng, 2015), consistent with its P^2^MR candidature. However, of critical importance to its potential utility as a therapeutic target, our study validates and extends this association beyond the non-synonymous variant to protein abundance. pQTL analysis identified two lead DNA variants in close proximity (<150kb) to the *FABP2* gene. Using the SNP rs17009129, P^2^MR finds a causal link between FABP2 concentration and CAD (p = 1.1×10^−4^; FDR <0.05; β_MR_ −0.11; se_MR_ 0.028; β_MR_ and se_MR_ units: log(OR)/standard deviation of residualised protein concentration) without significant heterogeneity (p = 0.24) which suggests shared causal genetic control. Furthermore, a second independent SNP (r^2^ <0.2; rs6857105) replicates this observation (MR p = 5.0×10-4; HEIDI p = 0.34; β_MR_ −0.17; se_MR_ 0.047). Both SNPs (rs17009129, and rs6857105) fell below genome-wide significance (p<5×10^−8^) in the full meta-analysis of van der Harst (van der Harst & Verweij, 2018) on CAD; however, FABP2 was flagged as potentially relevant by DEPICT, a prioritization tool. Consequently, this is the first time, to our knowledge, that variants associate with *FABP2* concentration have been shown robustly to causally contribute to CAD pathogenesis.

### Schizophrenia

By applying P^2^MR, we identified 3 proteins that were causally implicated in the pathogenesis of schizophrenia: (i) Tyrosine-protein phosphatase non-receptor type substrate 1 (SHPS1; *SIRPA*), (ii) Tumour necrosis factor receptor superfamily member 5 (*CD40*), and (iii) Low affinity immunoglobulin gamma Fc region receptor II-b (*FCGR2B*). The link between SHPS1 (rs4813319) and schizophrenia risk was subsequently replicated in the UK Biobank data (Methods; Table 1). The observed effect of SHSP1 on schizophrenia was not significantly heterogeneous in the results of the Schizophrenia Working Group of the Psychiatric Genomics Consortium (2014) (p = 0.53). Here we investigate *SHPS1* (approved symbol *SIRPA*), which encodes SHPS1, tyrosine-protein phosphatase non-receptor type substrate 1 and use *SHPS1* henceforth.

**Table 1:**
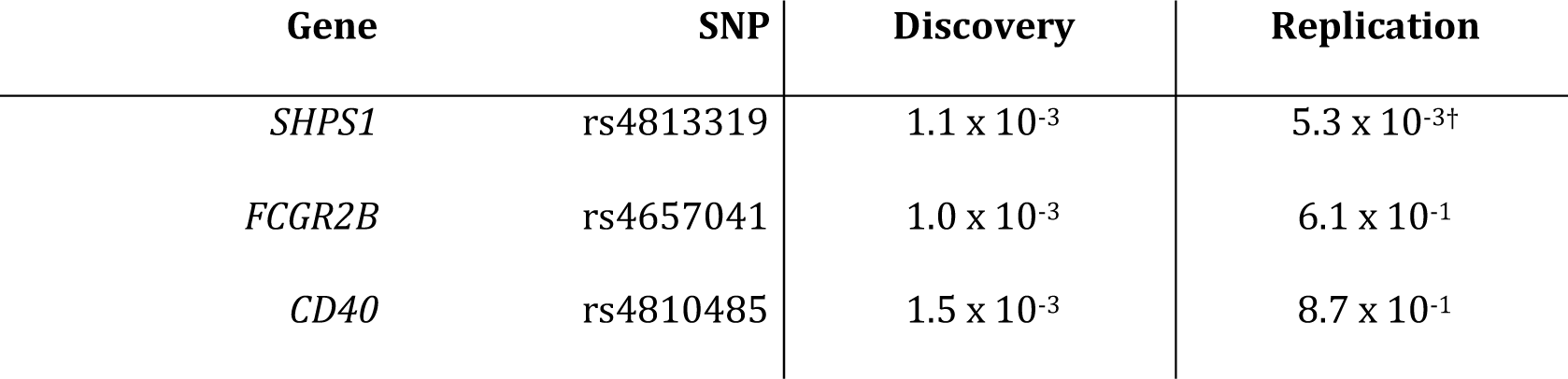
Replication of significant Mendelian Randomisation (FDR <0.05) protein-to-schizophrenia links in UK Biobank. Discovery: Mendelian randomisation p-value of protein level on schizophrenia risk, as estimated using data from the Psychiatric Genomics Consortium – PGC (Schizophrenia Working Group of the Psychiatric Genomics Consortium et al., 2014) obtained via Phenoscanner (Staley et al., 2016). Replication: The Mendelian Randomisation p-value of protein level on schizophrenia (combined risk of ‘F20-F29 Schizophrenia, schizotypal and delusional disorders’ and self-reported ‘schizophrenia’) in UK Biobank (Methods). MR p-values are significant (FDR <0.05) in the Discovery sample. † indicates significance of the replication study following multiple testing correction (Bonferroni).

| Gene | SNP | Discovery | Replication |
| --- | --- | --- | --- |
| *SHPS1* | rs4813319 | $1.1 \times 10^{-3}$ | $5.3 \times 10^{-3\dagger}$ |
| *FCGR2B* | rs4657041 | $1.0 \times 10^{-3}$ | $6.1 \times 10^{-1}$ |
| *CD40* | rs4810485 | $1.5 \times 10^{-3}$ | $8.7 \times 10^{-1}$ |

Interestingly, SHPS1 is highly expressed in the brain, especially in the neuropil (a dense network of axons, dendrites, and microglial cell processes) in the cerebral cortex (“SIRPA available from v18.proteinatlas.org,” 2018; “The Human Protein Atlas,” n.d.; Thul et al., 2017; Uhlén et al., 2015; Uhlen et al., 2017), and co-localises with CD47 at dendrite-axon contacts (Ohnishi et al., 2005). Mouse models in which the *SHPS1* gene is disrupted exhibit many nervous system abnormalities, such as reduced long term potentiation, abnormal synapse morphology and abnormal excitatory postsynaptic potential (MGI: 5558020 (“Mouse Genome Informatics (v6.13),” 2019; Toth et al., 2013)). Other mouse and rat models link CD47 to sensorimotor gating and social behaviour phenotypes (H. P. Chang, Lindberg, Wang, Huang, & Lee, 1999; Huang, Wang, Tang, & Lee, 1998; Koshimizu, Takao, Matozaki, Ohnishi, & Miyakawa, 2014; Ma, Kulesskaya, Võikar, & Tian, 2015; Ohnishi et al., 2010). In addition, SHPS1 has been shown to mediate activity-dependent synapse maturation (Toth et al., 2013) and may also have a role as a “don’t eat me” signal to microglia (Brown & Neher, 2014). Finally, SHPS1 levels tend to be lower in the dorsolateral prefrontal cortex of schizophrenia patients (Martins-de-Souza et al., 2009).

## Discussion

Proteome-by-phenome Mendelian Randomisation (P^2^MR) is an efficient method of identifying potential drug targets through integrating pQTL with myriad phenotypes. P^2^MR offers a data-driven approach to drug-discovery from population-level data. It quantifies the strength of evidence for causation, together with magnitude and direction of effect, for particular proteins in specific disease phenotypes. In addition, because MR using locally-acting pQTLs is more focussed than a genome-wide study, the burden of multiple testing is reduced dramatically, effectively reducing the sample-size required to declare a given effect significant.

P^2^MR has some inherent limitations that need to be considered when interpreting results. First, a true positive MR association in our analysis implies that any intervention to replicate the effect of a given genotype would alter the relevant phenotype. Nevertheless, this association is informative neither of the time interval, during development for example, nor the anatomical location in which an intervention would need to be delivered. Second, pleiotropic effects cannot be excluded entirely without (unachievable) quantification of every mediator. Third, the concentration of a protein in plasma could be an imperfect proxy for the effect of a drug targeting that protein at the level of a whole organism. Finally, plasma concentration does not necessarily reflect activity. For example, a variant may cause expression of high levels of an inactive form of a protein. Or, for proteins with both membrane-bound and unbound forms, the MR direction of effect observed from quantifying soluble protein abundance may not reflect that of membrane-bound protein. For many membrane-bound proteins, a soluble (often antagonistic) form exists that is commonly produced through alternative splicing or proteolytic cleavage of the membrane-bound form. For example, tocilizumab, an IL6 receptor antagonist, is used as a treatment of rheumatoid arthritis (L. J. Scott, 2017). The variant we use to instrument IL6R level, rs61812598, is in complete LD with the missense variant rs2228145 in the British sub-population of 1,000 Genomes (Sudmant et al., 2015; The 1000 Genomes Project Consortium, 2015) whose effects on proteolytic cleavage of the membrane-bound form and alternative splicing have been examined in detail (R. C. Ferreira et al., 2013). Carriers of the 358Ala allele at rs2228145 tend to have increased soluble IL6R but reduced membrane-bound IL6R in a number of immune cell types. Differences between the effects of soluble and membrane-bound forms of a protein may be wide-spread. For example, Dupilumab is a monoclonal antibody that targets IL4R, a key component of both IL4 and IL13 signalling. It is currently under investigation for the treatment of asthma and has shown promising results in both eosinophilic and non-eosinophilic asthma (Wenzel et al., 2016, 2013). Based on our results, we would have predicted that increased levels of IL4R result in a lower risk of asthma (Supplementary Table S3). This is in contrast to the direction-of-effect due to dupilumab administration. However, as with IL6R, IL4R has both a soluble and a membrane-bound form. Encouragingly, despite this, a relationship between dupilumab and asthma remains plausible – as evidenced by the 14 recently completed or ongoing clinical trials to assess the efficacy and safety of dupilumab in asthma (As of 26 Mar 2019, ClinicalTrials.gov Identifiers: NCT01312961, NCT01854047, NCT02134028, NCT02414854, NCT02528214, NCT02573233, NCT02948959, NCT03112577, NCT03387852, NCT03560466, NCT03620747, NCT03694158, NCT03782532, and NCT03884842).

P^2^MR provides an opportunity for studying the probable effects of specific proteins upon human diseases, such as schizophrenia, for which only imperfect model systems currently exist. Without a robust disease model, one must undertake studies in humans. However, there is little justification to undertake an adequately powered randomised control trial of a drug targeting a protein for which there is minimal evidence of a link between that protein and disease. P^2^MR does not suffer from such limitations.

P^2^MR highlights FABP2 as contributory to the pathogenesis of CAD and there are orthogonal lines of evidence to support this; notably: the non-synonymous mutation Ala54Thr (Yuan et al., 2015). In addition, given its interaction with PPAR-α and fenofibrate (Hughes et al., 2015) and strong expression in the gastrointestinal tract (“FABP2 available from v18.proteinatlas.org,” 2018; “The Human Protein Atlas,” n.d.; Thul et al., 2017; Uhlén et al., 2015; Uhlen et al., 2017), FABP2 represents a potential drug-target of the future.

Finally, as well as its utility in identifying potential therapeutic targets for drug development, P^2^MR allows for an assessment of potential off-target effects of existing pharmacological targets. For example, we predict an effect of IL4R modulation on eosinophil count and percentage. This is an association already realised in one of the phase II clinical trials investigating dupilumab in asthma: a rise in eosinophil count was observed for some patients, even leading to the withdrawal of one patient from the study (Wenzel et al., 2016, 2013).

## Conclusions

In summary, we have identified dozens of plausible causal links by conducting GWA of 249 proteins, followed by phenome-wide MR using replicated locally-acting pQTLs of 64 proteins: P^2^MR.

Using this approach, 54,144 protein-outcome links have been assessed and 509 significant (FDR <0.05) links identified: including anthropometric measures, haematological parameters, as well as diseases. Opportunities to discover larger sets of plausible causal links will increase as study sizes and pQTL numbers grow. Indeed, whole-proteome versus Biobank GWA Atlas studies will likely become feasible as pQTL measurement technologies mature.

## Methods

### Cohort description

From the islands of Orkney (Scotland) and Vis (Croatia) respectively, the ORCADES (McQuillan et al., 2008) and CROATIA-Vis (Campbell et al., 2007; Rudan et al., 2009) studies are of two isolated population cohorts that are both genotyped and richly phenotyped.

The Orkney Complex Disease Study (ORCADES) is a family-based, cross-sectional study that seeks to identify genetic factors influencing cardiovascular and other disease risk in the isolated archipelago of the Orkney Isles in northern Scotland (McQuillan et al., 2008). Genetic diversity in this population is decreased compared to Mainland Scotland, consistent with the high levels of endogamy historically. 2,078 participants aged 16-100 years were recruited between 2005 and 2011, most having three or four grandparents from Orkney, the remainder with two Orcadian grandparents. Fasting blood samples were collected and many health-related phenotypes and environmental exposures were measured in each individual. All participants gave written informed consent and the study was approved by Research Ethics Committees in Orkney and Aberdeen (North of Scotland REC).

The CROATIA-Vis study includes 1,008 Croatians, aged 18-93 years, who were recruited from the villages of Vis and Komiza on the Dalmatian island of Vis during spring of 2003 and 2004. All participants were volunteers and gave informed consent. They underwent a medical examination and interview, led by research teams from the Institute for Anthropological Research and the Andrija Stampar School of Public Health, (Zagreb, Croatia). All subjects visited the clinical research centre in the region, where they were examined in person and where fasting blood was drawn and stored for future analyses. Many biochemical and physiological measurements were performed, and questionnaires of medical history as well as lifestyle and environmental exposures were collected. The study received approval from the relevant ethics committees in Scotland and Croatia (REC reference: 11/AL/0222) and complied with the tenets of the Declaration of Helsinki.

### Genotyping

Chromosomes and positions reported in this paper are from GRCh37 throughout. Genotyping of the ORCADES cohort was performed on the Illumina Human Hap 300v2, Illumina Omni Express, and Illumina Omni 1 arrays; that of the CROATIA-Vis cohort used the Illumina HumanHap300v1 array.

The genotyping array data were subject to the following quality control thresholds: genotype call-rate 0.98, per-individual call-rate 0.97, failed Hardy-Weinberg test at p-value *<* 1 × 10^−6^, and minor allele frequency 0.01; genomic relationship matrix and principal components were calculated using GenABEL (1.8-0) (Aulchenko, Ripke, Isaacs, & van Duijn, 2007) and PLINK v1.90 (C. C. Chang et al., 2015; Purcell, 2017).

Assessment for ancestry outliers was performed by anchored PCA analysis when compared to all non-European populations from the 1,000 Genomes project (Sudmant et al., 2015; The 1000 Genomes Project Consortium, 2015). Individuals with a mean-squared distance of >10% in the first two principal components were removed. Genotypes were phased using Shapeit v2.r873 and duoHMM (O’Connell et al., 2014) and imputed to the HRC.r1-1 reference panel (The Haplotype Reference Consortium et al., 2016). 278,618 markers (Hap300) and 599,638 markers (Omni) were used for the imputation in ORCADES, and 272,930 markers for CROATIA-Vis.

### Proteomics

Plasma abundance of 249 proteins was measured in two European cohorts using Olink Proseek Multiplex CVD2, CVD3, and INF panels. All proteomics measurements were obtained from fasting EDTA plasma samples. Following quality control, there were 971 individuals in ORCADES, and 887 individuals in CROATIA-Vis, who had genotype and proteomic data from Olink CVD2, 993 and 899 from Olink CVD3, and 982 and 894 from Olink INF. The Olink Proseek Multiplex method uses a matched pair of antibodies for each protein, linked to paired oligonucleotides. Binding of the antibodies to the protein brings the oligonucleotides into close proximity and permits hybridization. Following binding and extension, these oligonucleotides form the basis of a quantitative PCR reaction that allows relative quantification of the initial protein concentration (Assarsson et al., 2014). Olink panels include internal and external controls on each plate: two controls of the immunoassay (two non-human proteins), one control of oligonucleotide extension (an antibody linked to two matched oligonucleotides for immediate proximity, independent of antigen binding) and one control of hybridized oligonucleotide detection (a pre-made synthetic double stranded template), as well as an external, between-plate, control (“Olink,” n.d.).

Prior to analysis, we excluded proteins with fewer than 200 samples with measurements above the limit of detection of the assay. Of the 268 unique proteins reported by Olink, 253 passed this threshold in ORCADES, and 252 in CROATIA-Vis, with an intersect of 251 proteins. Protein values were inverse-normal rank-transformed prior to subsequent analysis.

The subunits of IL27 are not distinguished in Olink’s annotation (Q14213, *EBI3*; and Q8NEV9, *IL27*). However, it has only one significant locus, local to the *EBI3* gene (lead variant, rs60160662, is within 16kb). Therefore, *EBI3* (Q14213) was selected as representative for this protein when discussing pQTL location (local/distal) so as to avoid double counting.

Two proteins, CCL20 and BDNF, have been removed at the request of Olink.

### Genome-wide association of protein levels

Genome-wide association of these proteins was performed using autosomes only. Analyses were performed in three-stages. (1) a linear regression model was used to account for participant age, sex, genotyping array (ORCADES only), proteomics plate, proteomics plate row, proteomics plate column, length of sample storage, season of venepuncture (ORCADES only), and the first 10 principal components of the genomic relationship matrix. Genotyping array and season of venepuncture are invariant in CROATIA-Vis and therefore were not included in the model. (2) Residuals from this model were corrected for relatedness, using GenABELs (Aulchenko et al., 2007) polygenic function and the genomic relationship matrix, to produce GRAMMAR+ residuals. Outlying GRAMMAR+ residuals (absolute z-score >4) were removed and the remainder rank-based inverse-normal transformed. (3) Genome-wide association testing was performed using REGSCAN v0.5 (Haller, Kals, Esko, Magi, & Fischer, 2013).

### Reported pQTLs

Genome-wide association results were clumped by linkage disequilibrium using PLINK v1.90 (C. C. Chang et al., 2015; Purcell, 2017). Biallelic variants within ±5Mb and *r*^2^ >0.2 to the lead variant (smallest p-value at the locus) were clumped together, and the lead variant is presented. *r*^2^ was derived from all European populations in 1,000 Genomes (Sudmant et al., 2015; The 1000 Genomes Project Consortium, 2015).

### Mendelian Randomisation

In the context of P^2^MR, a DNA variant (a single nucleotide polymorphism in this case) that influences plasma protein level is described as an ‘instrumental variable’, the protein as the ‘exposure variable’, and the outcome phenotype as the ‘outcome variable’.

A DNA variant was considered to be a potentially valid instrumental variable if it met the following criteria:

1. Minor allele frequency >1% in both ORCADES and CROATIA-Vis cohorts.
2. An imputation info score (SNPTEST v2) of >0.95 in both ORCADES and CROATIA-Vis.
3. Located within ±150kb of the gene coding for the protein (start and end coordinates of the gene as defined by Ensembl GRCh37 (Zerbino et al., 2018)).

DNA variant selection was performed using the discovery (CROATIA-Vis) cohort. Replication was defined based on a Bonferroni correction for the number of genome-wide significant lead variants selected in the discovery cohort (CROATIA-Vis). In order to avoid a ‘winner’s curse’, replicated genome-wide association effect sizes and standard errors from the replication cohort (ORCADES) were used for MR.

We perform MR as a ratio of expectations, using up to second-order partial derivatives of the Taylor series expansion for effect size estimates, and up to first-order for standard errors (Delta method) (Lynch & Walsh, 1998):

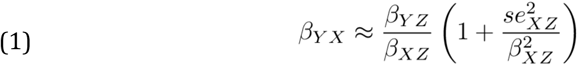

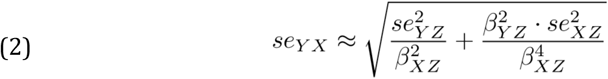

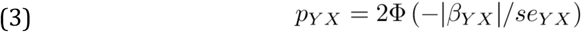

where *β*_*ij*_ is the causal effect of *j* on *i, se*_*ij*_ is the standard error of the causal effect estimate of *j* on *i*; subscript *X* is the exposure, *Y* the outcome trait, and *Z* the instrumental variable. F is the cumulative density function of the standard normal distribution.

### DNA variant to trait association: GeneAtlas

All outcome GWA (778 traits) from GeneAtlas (Canela-Xandri et al., 2017) were included. For each protein, the lead (lowest DNA variant-protein association p-value in the discovery cohort) biallelic (Phase 3, 1,000 Genomes (Sudmant et al., 2015; The 1000 Genomes Project Consortium, 2015)) variant meeting the criteria above and an imputation info score >0.95 in UK Biobank, was selected for each protein, and MR performed. An FDR of <0.05 was considered to be significant.

### DNA variant to trait association: Phenoscanner

Phenoscanner (“PhenoScanner,” 2018; Staley et al., 2016) was used to highlight existing GWA studies for inclusion. For each protein, the lead (lowest DNA variant-protein association p-value in the discovery cohort) biallelic (1,000 Genomes (Sudmant et al., 2015; The 1000 Genomes Project Consortium, 2015)) meeting the criteria above was selected. rs545634 was not found in the Phenoscanner database and was therefore replaced with the second most significant variant meeting the above criteria: chr1:15849003. Phenoscanner was run with the following options: Catalogue: ‘Diseases & Traits’, p-value cut-off: ‘1’, Proxies: ‘None’, Build ‘37’. Results from 20 additional studies were obtained, corresponding to 68 outcomes. The results from those studies that returned a value for all input variants were kept and MR performed. An FDR of <0.05 was considered to be significant.

### HEIDI

Heterogeneity in dependent instruments (HEIDI) analysis (Zhu et al., 2016), is a method of testing whether the MR estimates obtained using variants in linkage disequilibrium with the lead variant are consistent with a single causal variant or multiple causal variant at a given locus (Figure 1D). HEIDI analysis was performed using software provided at https://cnsgenomics.com/software/smr/ [accessed 28/08/2018; v0.710]. We created a bespoke BESD format file containing the pQTL data from ORCADES for assessment as the exposure. Biallelic variants from the 1,000 Genomes (Sudmant et al., 2015; The 1000 Genomes Project Consortium, 2015) (European populations: CEU, FIN, GBR, IBS, and TSI) were used as the linkage disequilibrium reference. We used the default ‘cis-window’ of 2000kb, and a maximum number of variants of 20 (as this is now the default value for the software: based on unpublished power calculations by the authors of HEIDI and noted on their website).

We performed HEIDI analysis of all exposure-outcome links that were found to be significant (FDR <0.05) using outcomes from UK Biobank (n=271), as well as those links found to be MR significant (FDR <0.05) with CAD from the meta-analysis of van der Harst (van der Harst & Verweij, 2018), and for SHPS1 and schizophrenia (Schizophrenia Working Group of the Psychiatric Genomics Consortium et al., 2014).

We applied the following filters for variants to be included in the analysis: minor allele frequency MAF *>* 0.01 and, in the GeneAtlas and ORCADES data, an imputation info score of >0.95.

### Schizophrenia GWA study replication

In the initial analysis of Canela-Xandri et al. (2017), schizophrenia was included as ‘F20 Schizophrenia’, and nested in ‘F20-F29 Schizophrenia, schizotypal and delusional disorders’. There were 920 cases in ‘F20-F29 Schizophrenia, schizotypal and delusional disorders’ and 509 in ‘F20 Schizophrenia’. Due to the near doubling of the sample size, replication was attempted in the parent category: ‘F20-F29 Schizophrenia, schizotypal and delusional disorders’. Using a Bonferroni correction, none of these links replicated. However, due to the severe contraction of the number of cases present in the sample – 35,476 cases and 46,839 controls to 920 cases and 407,535 controls – there was a significant risk of false negative results. In order to address this, we re-analysed the UK Biobank data including ‘F20-F29 Schizophrenia, schizotypal and delusional disorders’ and self-reported ‘schizophrenia’ as a single outcome in a more permissive set of individuals: individuals self-reporting their ethnicity as ‘White’ and clustering as a group based on the first two genomic principal components (Canela-Xandri, Rawlik, & Tenesa, 2018). This increased the number of cases and controls to 1,241 cases and 451,023 controls.

## Supporting information

Table S2 - columns

Table S3 - columns

Table S4 - columns

## Acknowledgements

- A debt of gratitude is owed to all the participants in all cohorts used, without whom this work would not have been possible.
- This research has been conducted using the UK Biobank Resource under project 788.
- Funding
  - ADB would like to acknowledge funding from the Wellcome PhD training fellowship for clinicians (204979/Z/16/Z), the Edinburgh Clinical Academic Track (ECAT) programme.
  - TB, YZ, CA, PN, JFW, VV, CHay, CPP and CHal are supported by MRC University Unit Programme Grants to the Human Genetics Unit (MC_PC_U127592696, MC_UU_12008/1, MC_UU_00007/10 and MC_UU_00007/15)
  - AT, OC-X and KR acknowledge funding from the MRC (MR/R025851/1, MR/N003179/1).
  - CHal, JKB, AT, and KR acknowledge funding from BBSRC Institute Strategic Programme grants to the Roslin Institute (BBS/E/D/30002275, BBS/E/D/30002276, BBS/E/D/10002071, BBS/E/D/20002172, BBS/E/D/20002174).
  - PKJ would like to acknowledge funding from the Axa research fund.
  - JKB acknowledges funding support from a Wellcome-Beit Prize Intermediate Clinical Fellowship (103258/Z/13/Z,A), and the UK Intensive Care Foundation.
- The Orkney Complex Disease Study (ORCADES) was supported by the Chief Scientist Office of the Scottish Government (CZB/4/276, CZB/4/710), a Royal Society URF to JFW, the MRC Human Genetics Unit quinquennial programme “QTL in Health and Disease”, Arthritis Research UK and the European Union framework program 6 EUROSPAN project (contract no. LSHG-CT-2006-018947). DNA extractions were performed at the Wellcome Trust Clinical Research Facility in Edinburgh. We would like to acknowledge the invaluable contributions of the research nurses in Orkney, the administrative team in Edinburgh and the people of Orkney.
- The CROATIA-Vis study was funded by grants from the Medical Research Council (UK) and Republic of Croatia Ministry of Science, Education and Sports research grants. (108-1080315-0302). We would like to acknowledge the staff of several institutions in Croatia that supported the field work, including but not limited to The University of Split and Zagreb Medical Schools, the Institute for Anthropological Research in Zagreb and Croatian Institute for Public Health. Genotyping was performed in the Genetics Core of the Clinical Research Facility, University of Edinburgh.

## Supplementary Materials

- Table S1. Additional studies identified using Phenoscanner: additional_studies.tsv.
- Table S2. Complete list of pQTLs (linkage disequilibrium clumped): indep_pqtl.tsv.
- Table S3. Mendelian Randomisation results from UK Biobank: df_ukbb_heidi.tsv.
- Table S4. Mendelian Randomisation results from studies identified using Phenoscanner: df_phenoscanner.tsv.
- Figure S1. Significant (FDR <0.05) P2MR protein-outcome causal inferences: haematology count subset.
- Figure S2. Significant (FDR <0.05) P2MR protein-outcome causal inferences: haematology percentage subset.
- Figure S3. Significant (FDR <0.05) P2MR protein-outcome causal inferences: haematology (non-count, non-percentage) subset.
- Figure S4. Significant (FDR <0.05) P2MR protein-outcome causal inferences: anthropometric measurements subset.

**Supplementary Figure S1.**
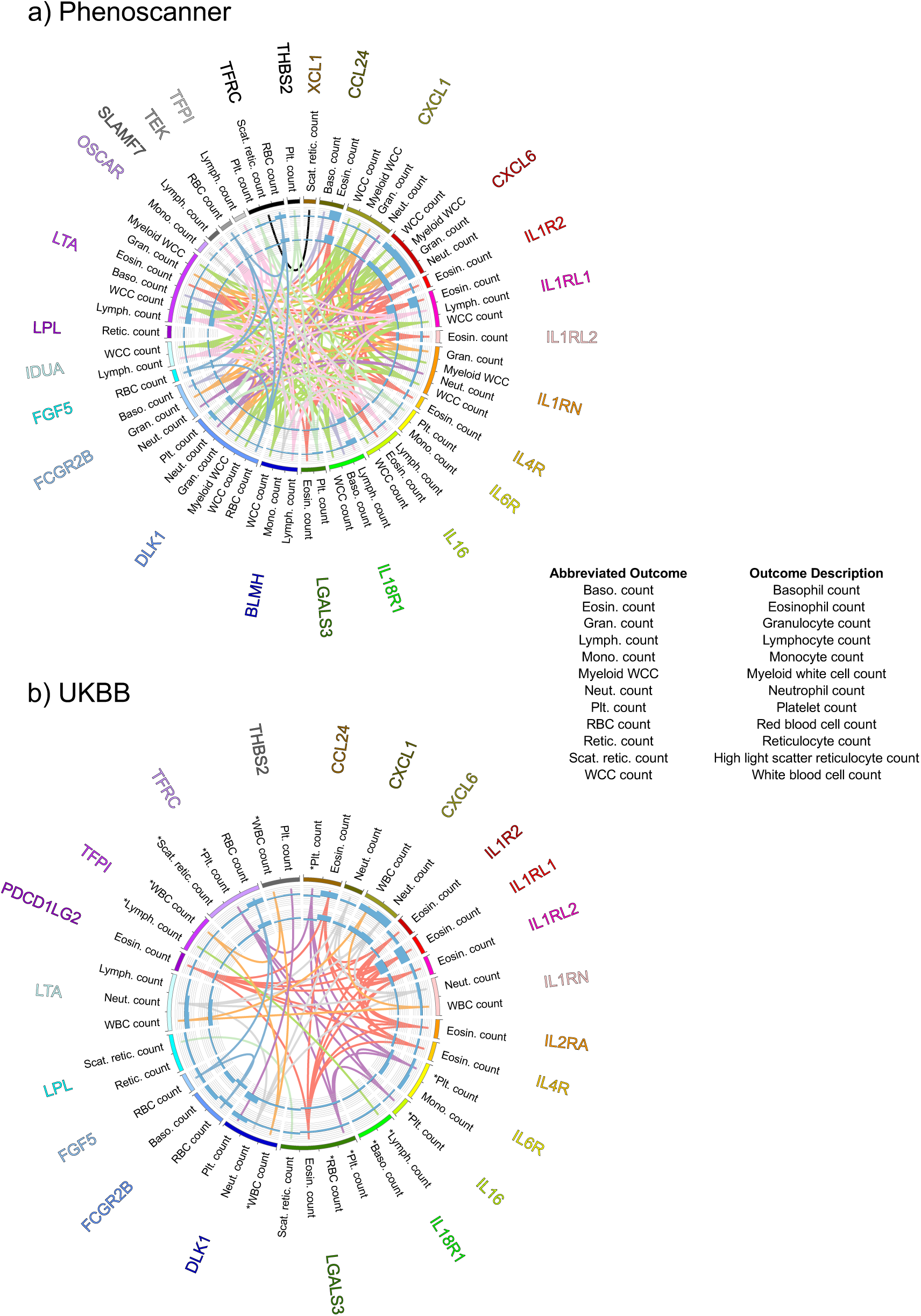
Significant (FDR <0.05) P^2^MR protein-outcome causal inferences: haematology count subset. a) Phenoscanner (Staley et al., 2016): P^2^MR significant protein-haematology count outcome causal inferences for 20 Phenoscanner studies. b) GeneAtlas (Canela-Xandri et al., 2017): MR significant protein-haematology count outcome causal inferences for UK Biobank data. Asterisks indicate P^2^MR estimates that are not significantly heterogeneous (HEIDI, Main Text (Zhu et al., 2016)). Key as for Figure 2: Reading from the outside in: protein (exposure; HGNC symbol); haematology count outcome; key colour; bar chart of the signed (beta/standard error)^2^ value of the MR estimate (using pQTL data from the discovery cohort; Methods); and bar chart of the signed (beta/standard error)^2^ value of the MR estimate (using pQTL data from the replication cohort; Methods). Central chords join identical outcomes. Identically coloured chords indicate similar outcome groups.

**Supplementary Figure S2.**
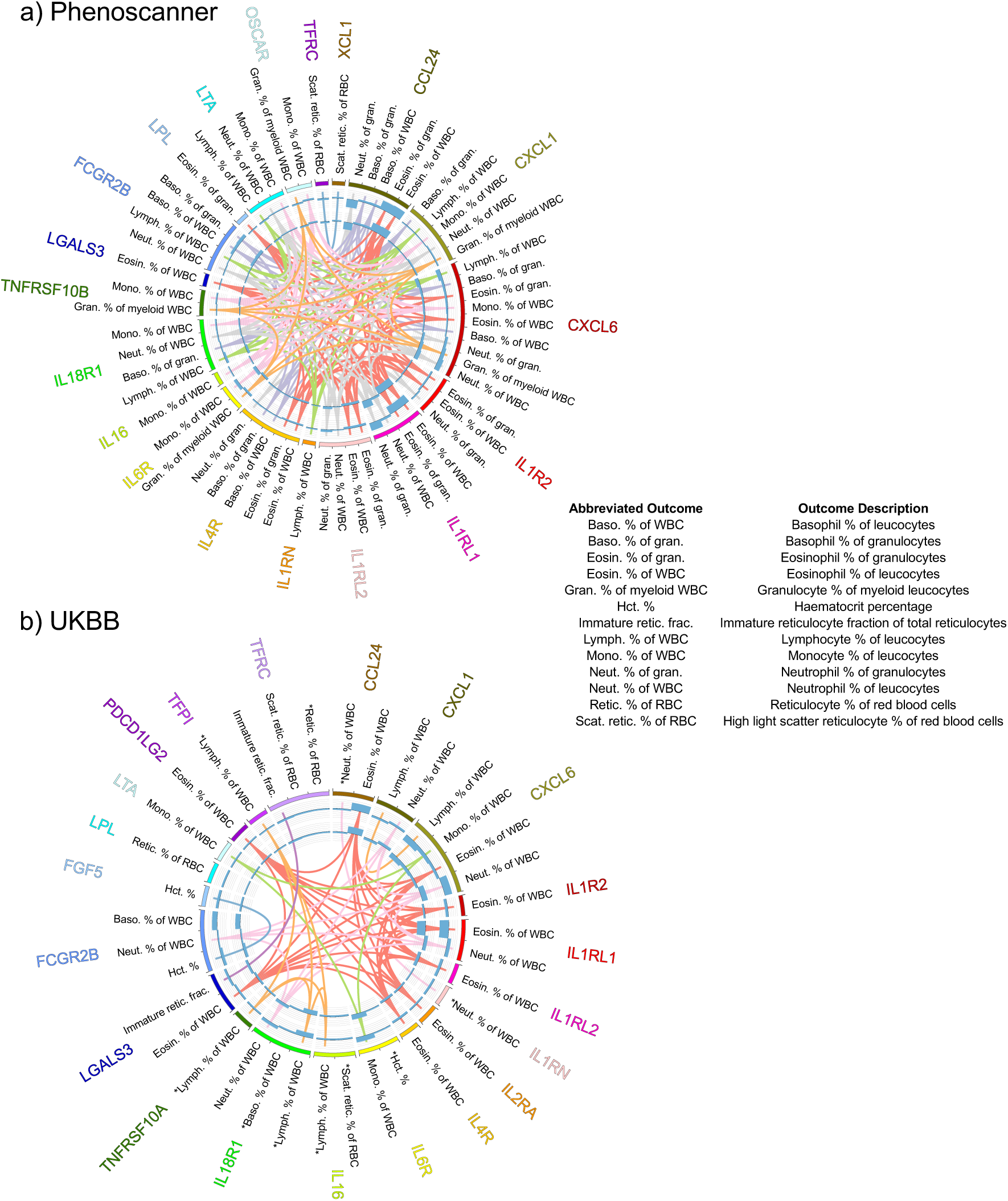
Significant (FDR <0.05) P^2^MR protein-outcome causal inferences: haematology percentage subset. a) Phenoscanner (Staley et al., 2016): P^2^MR significant protein-haematology percentage outcome causal inferences for 20 Phenoscanner studies. b) GeneAtlas (Canela-Xandri et al., 2017): MR significant protein-haematology percentage outcome causal inferences for UK Biobank data. Asterisks indicate P^2^MR estimates that are not significantly heterogeneous (HEIDI, Main Text (Zhu et al., 2016)). Key as for Figure 2: Reading from the outside in: protein (exposure; HGNC symbol); haematology percentage outcome; key colour; bar chart of the signed (beta/standard error)^2^ value of the MR estimate (using pQTL data from the discovery cohort; Methods); and bar chart of the signed (beta/standard error)^2^ value of the MR estimate (using pQTL data from the replication cohort; Methods). Central chords join identical outcomes. Identically coloured chords indicate similar outcome groups.

**Supplementary Figure S3.**
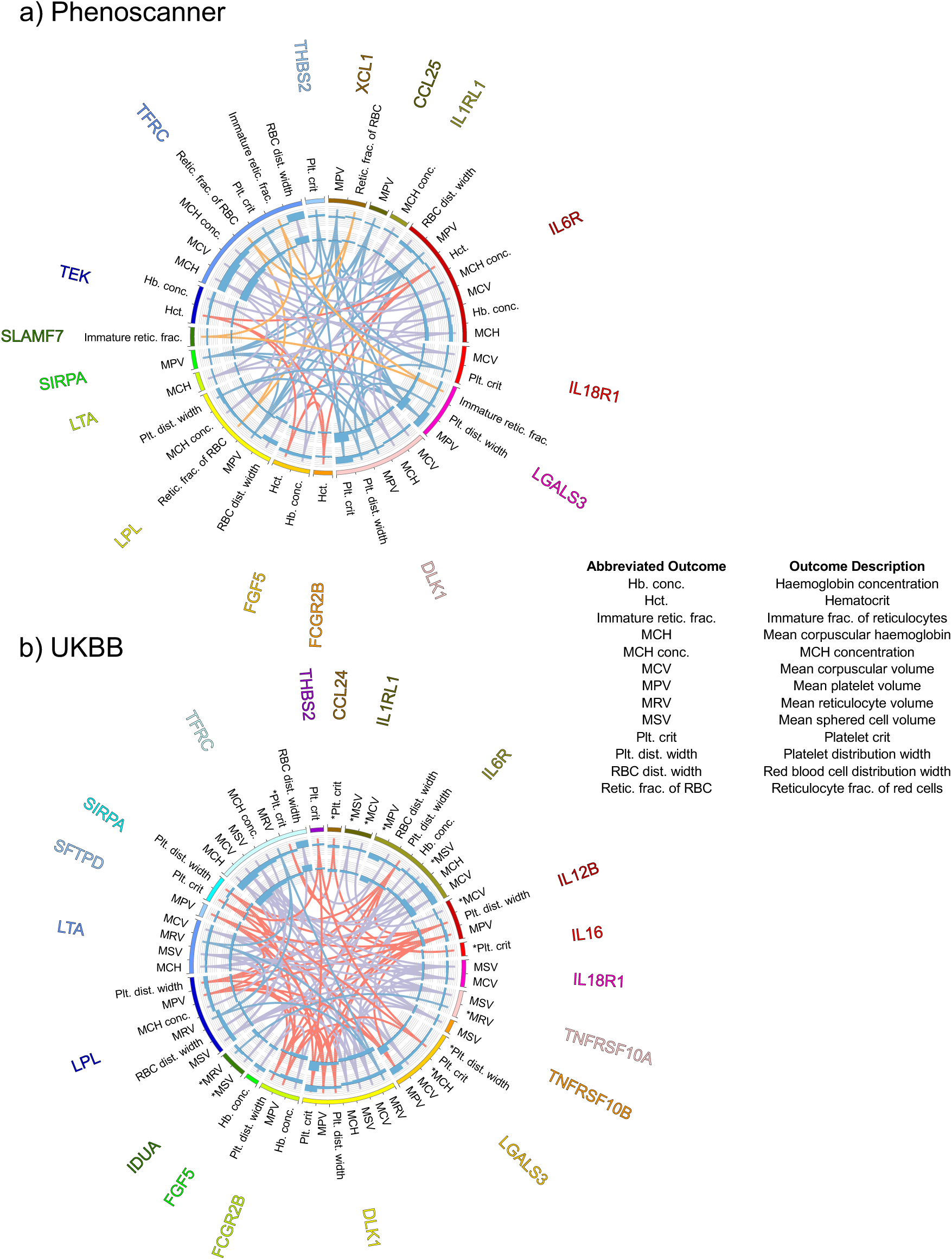
Significant (FDR <0.05) P^2^MR protein-outcome causal inferences: haematology (non-count, non-percentage) subset. a) Phenoscanner (Staley et al., 2016): P^2^MR significant protein-haematology outcome causal inferences for 20 Phenoscanner studies. b) GeneAtlas (Canela-Xandri et al., 2017): MR significant protein-haematology outcome causal inferences for UK Biobank data. Asterisks indicate P^2^MR estimates that are not significantly heterogeneous (HEIDI, Main Text (Zhu et al., 2016)). Key as for Figure 2: Reading from the outside in: protein (exposure; HGNC symbol); haematology outcome; key colour; bar chart of the signed (beta/standard error)^2^ value of the MR estimate (using pQTL data from the discovery cohort; Methods); and bar chart of the signed (beta/standard error)^2^ value of the MR estimate (using pQTL data from the replication cohort; Methods). Central chords join identical outcomes. Identically coloured chords indicate similar outcome groups.

**Supplementary Figure S4.**
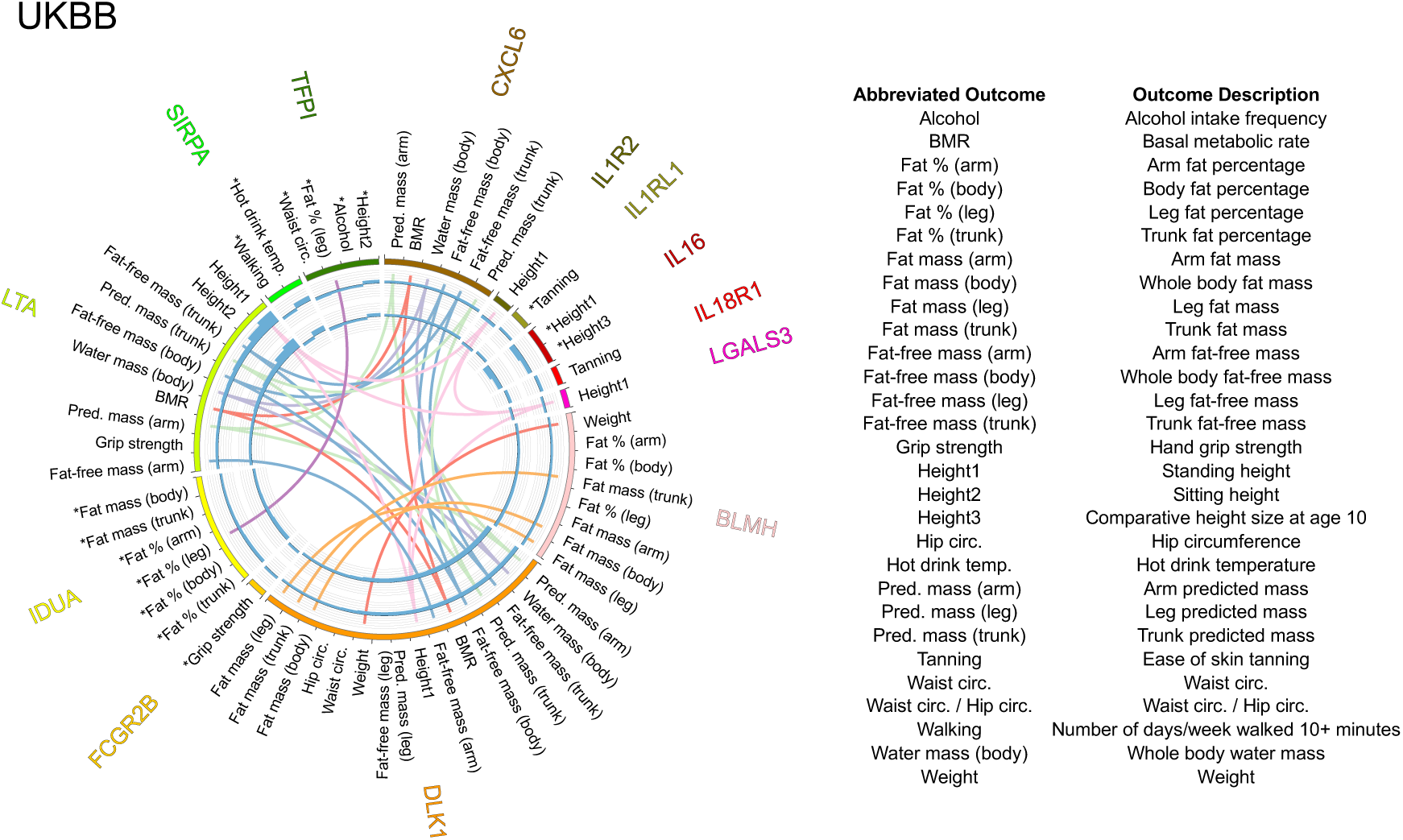
Significant (FDR <0.05) P^2^MR protein-outcome causal inferences: anthropometric measurements subset. GeneAtlas (Canela-Xandri et al., 2017): MR significant protein-anthropometric measurements outcome causal inferences for UK Biobank data. Asterisks indicate P^2^MR estimates that are not significantly heterogeneous (HEIDI, Main Text (Zhu et al., 2016)). Key as for Figure 2: Reading from the outside in: protein (exposure; HGNC symbol); anthropometric measurements outcome; key colour; bar chart of the signed (beta/standard error)^2^ value of the MR estimate (using pQTL data from the discovery cohort; Methods); and bar chart of the signed (beta/standard error)^2^ value of the MR estimate (using pQTL data from the replication cohort; Methods). Central chords join identical outcomes. Identically coloured chords indicate similar outcome groups.

